# All-trans retinoic acid (ATRA) reduces proliferative capacity and Brachyury levels in chordoma cells

**DOI:** 10.1101/2020.01.09.897314

**Authors:** Helena Robinson, Ramsay J. McFarlane, Jane A. Wakeman

## Abstract

Chordoma is a rare bone cancer for which there are no approved drugs. Surgery is the principle treatment but complete resection can be challenging due to the location of the tumours in the spine and therefore finding an effective drug treatment is a pressing unmet clinical need. A major recent study identified the transcription factor Brachyury as the primary vulnerability and drug target in chordoma. Previously, all-trans retinoic acid (ATRA) has been shown to negatively influence expression of the Brachyury gene, *TBXT*. Here we extend this finding and demonstrate that ATRA lowers Brachyury protein levels in chordoma cells and reduces proliferation of the chordoma cell line U-CH1 as well as causing a morphological change commensurate with differentiation. ATRA is available as a generic drug and is the first line treatment for acute promyelocytic leukaemia (APL). This study implies ATRA could have therapeutic value if repurposed for chordoma.

## Introduction

Chordoma is a rare bone cancer with an annual incidence of 1 in 1,000,000 people and a poor prognosis. It occurs mostly in the spine and whilst surgical resection is currently the most effective treatment this is difficult due to proximity to important structures. Chordomas are largely refractory to current chemotherapy, therefore, there is a pressing unmet clinical need for effective treatments (Stacchiotti & Sommer, 2015). Recently, the transcription factor Brachyury was identified as a primary drug target in chordoma (Sharifnia et al., 2019).

All-trans retinoic acid (ATRA) is available as a generic drug and is used as a differentiation agent to treat acute promyelocytic leukaemia (APL; Fey & Buske, 2013). U-CH1 chordoma cells treated with ATRA have reduced levels of *TBXT* (Brachyury gene) mRNA and reduced proliferative capacity (Aydemir et al., 2012). This study extends and validates these findings offering further insight into the mechanisms of ATRA as a potential chordoma therapeutic agent.

## Objective

The finding that ATRA reduces levels of *TBXT* (Brachyury) mRNA and reduces proliferative capacity opens three important questions. Firstly, does the reduction in mRNA levels feed through to measurable reduction in Brachyury protein? Secondly, is this effect of ATRA specific to the previously tested chordoma cell line, or is it a universal feature of chordoma cells? Finally, is the reduced proliferative capacity induced by ATRA in chordoma cells, which correlates to Brachyury loss, accompanied by the morphological changes observed following specific siRNA-mediated Brachyury depletion? The objectives of this current study are to address these outstanding and important questions.

## Methods and materials

### Cell culture and drug treatment

U-CH1 and JHC7 were obtained from ATCC and cultured as per ATCC guidelines. For all experiments described, 7.75 ×10^4^ cells were seeded in each well of a 6 well plate. ATRA (Abcam) and DMSO (Sigma Aldrich) were added to the concentrations specified. Drug and media were refreshed every 3 days. Cell counting was performed using a TC20 cell counter (Biorad). Cells were imaged using an Evos Core microscope (AMG).

### Protein extraction and western blot

Protein was extracted using M-PER buffer (ThermoFisher Scientific) according to the manufacturer’s instructions. Samples were denatured in Bolt LDS Sample buffer and Reducing Agent and run on an SDS-polyacrylamide pre cast gel (Bolt 4-12% Bis-Tris Plus, Life Technologies). Samples were transferred onto PVDF membrane in Towbin buffer (10% methanol) and the membrane was blocked in blocking solution (10% skimmed milk in PBS 0.1% Tween20). The membrane was incubated with primary antibody in blocking solution overnight and the membrane was incubated with secondary antibody at room temperature for 2 hours. Primary antibodies used were anti-Brachyury (Abcam, ab209665) and anti-GapDH (Santa Cruz, sc365062). Secondary (HRP linked) antibodies used were: anti-rabbit IgG (CST, 7074S), anti-mouse IgG (CST, 7076S).

## Results

### Treatment of chordoma cells with ATRA reduces Brachyury levels

JHC7 and U-CH1 chordoma cells were treated with 10 or 20 μM ATRA. ATRA caused a reduction in Brachyury levels in both lines, although this was less pronounced in JHC7 (Fig.1).

**Figure 1.**
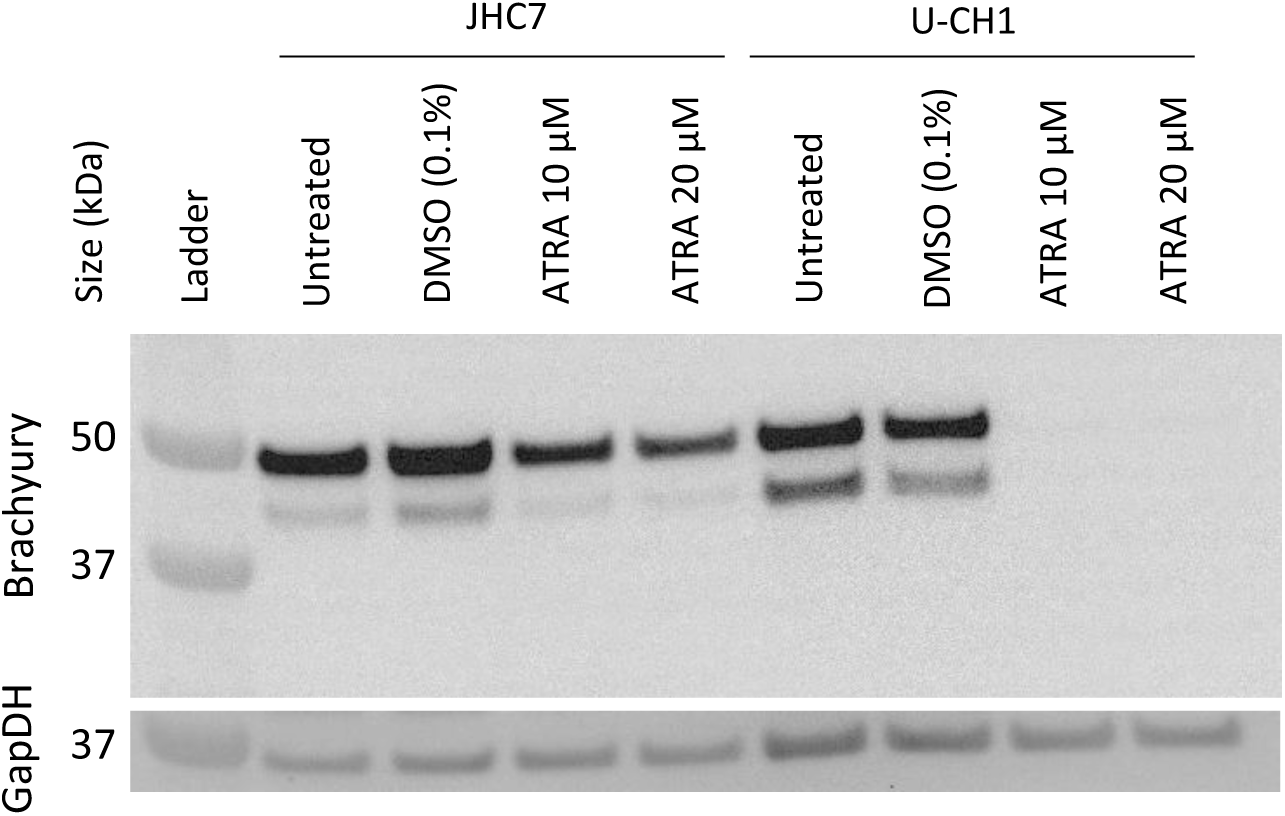
Western blot showing Brachyury levels in two chordoma cell lines following no treatment or treatment with DMSO or ATRA. U-CH1 cells were treated for 6 days and JHC7 cells were treated for 9 days. The same membrane was reprobed with anti-GAPDH antibody as a loading control. The blots were imaged using a Biorad Chemidoc. Colorimetric and chemiluminscent images were combined to show ladder and protein detection in the same image. This western blot is representative of 2 independent repeats.

### Treatment of chordoma cells with ATRA causes reduced proliferative capacity and morphological differentiation

U-CH1 cells treated with ATRA have reduced proliferative capacity (Aydimer et al., 2012). We treated U-CH1 cells with 20 μM ATRA and this resulted in the cell culture failing to increase cell numbers, assumed to be a proliferative inhibition, thus validating the original study (Fig. 2). To extend this we assessed whether there were any morphological changes apparent commensurate with those previously observed following specific siRNA-mediated Brachyury depletion (Hsu et al., 2011). ATRA treatment resulted in a loss of the physaliferous phenotype and the cells became more elongated and branching (Fig. 3).

**Figure 2.**
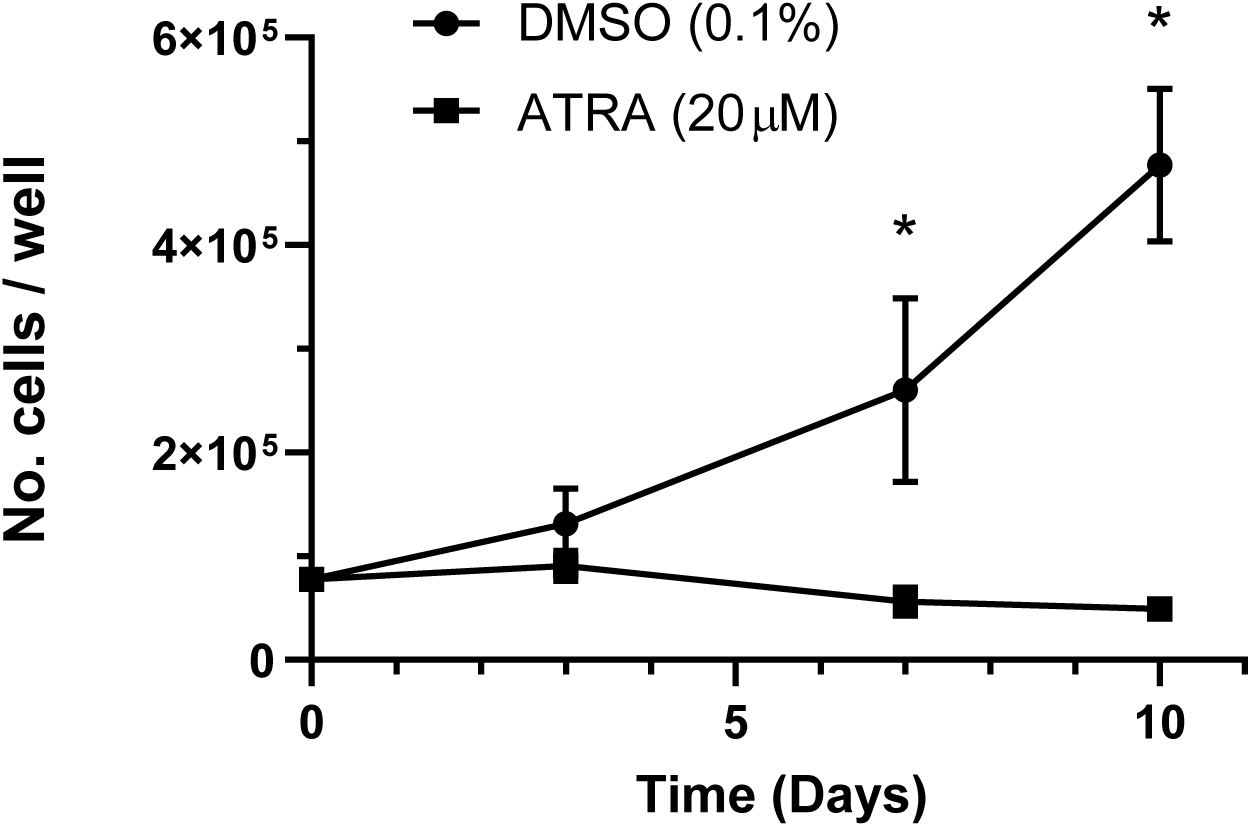
Cell number plot for U-CH1 with and without ATRA treatment. Values shown are the mean of two independent repeats. Error bars show standard error of the mean. Asterisks denote statistical significance < p = 0.05 (multiple t-tests using the Holm-Sidak method, equal variance, no. t tests = 4).

**Figure 3.**
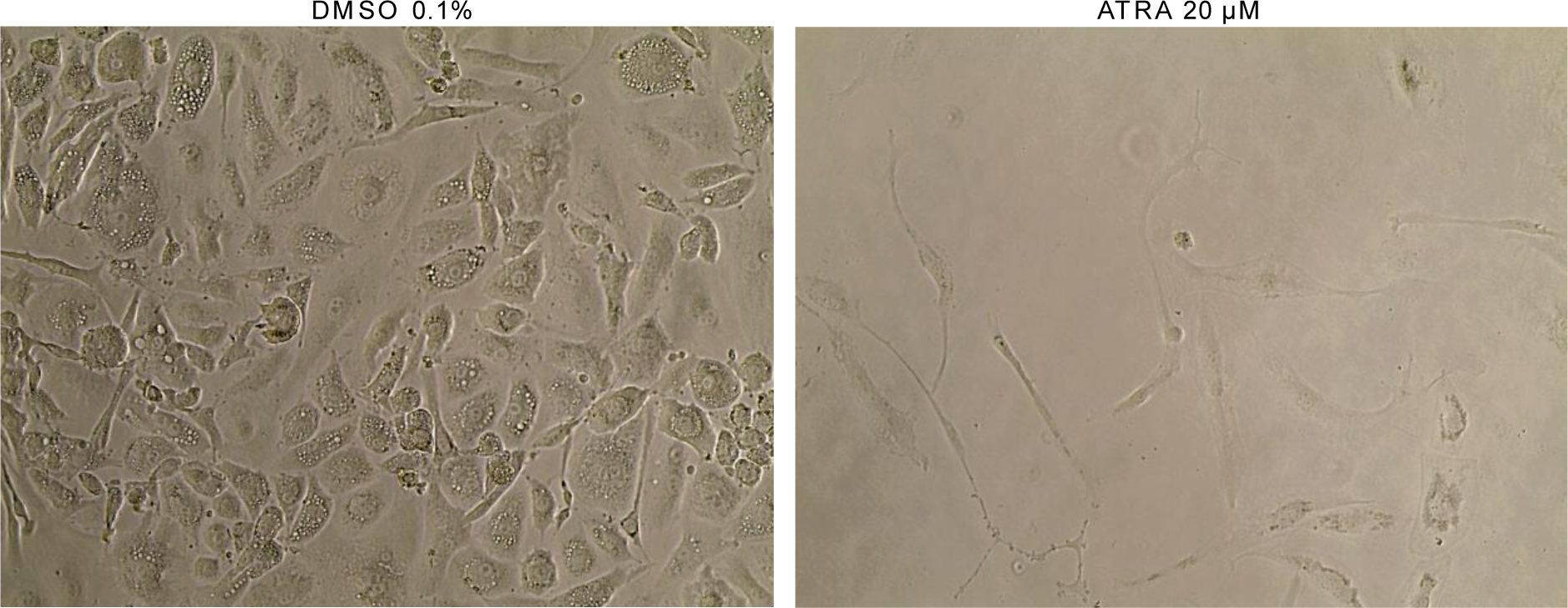
Brightfield images of U-CH1 cells after treatment with either DMSO or ATRA 20 μM for 10 days, illustrating the morphology changes observed for the whole cell population. The pictures are representative of two independent repeats. Images captured using x20 objective.

## Discussion

Brachyury has emerged as a target for the treatment of chordoma. The finding that ATRA results in the reduction of *TBXT* (Brachyury) mRNA brings ATRA into consideration as a repurposed therapeutic intervention. Here we have added important additional insight into the relationship between the ATRA response pathway and Brachyury in chordoma cells. Our findings support a model in which ATRA activates differentiation, which involves the shutdown of Brachyury activity. Indeed, such a model is supported by work in zebrafish notochord development, where ATRA treatment rapidly reduces levels of transcripts from the zebrafish orthologue gene, *ntl* (Martin & Kimelman, 2010).

## Conclusions

This study demonstrates that ATRA treatment lowers Brachyury levels, consistent with mRNA studies (Aydemir et al., 2012). The extent to which ATRA influences Brachyury levels varies between chordoma cells types, so targeting Brachyury up-stream regulatory pathways with ATRA might not prove to be universally effective. It remains unclear if Brachyury is directly regulated by an ATRA response pathway, but we can conclude that response to ATRA cellular changes are highly similar to those observed for specific Brachyury depletion. Thus, repurposing of ATRA to target a Brachyury activating pathway is an important consideration.

## Author contributions, data availability statement, conflicts of interest and financial declaration

HR and JAW conceived and designed the study. HR conducted data gathering. HR performed statistical analyses. HR, JAW and RJM wrote the article.

The data that support the findings of this study are openly available in the Open Science Framework at http://doi.org/10.17605/OSF.IO/C2V6E, reference number [C2V6E].

